# *Cryptocercus* genomes expand knowledge of adaptations to xylophagy and termite sociality

**DOI:** 10.1101/2025.01.24.634722

**Authors:** Alun R. C. Jones, Alina A. Mikhailova, Cédric Aumont, Juliette Berger, Erich Bornberg-Bauer, Cong Liu, Shulin He, Zongqing Wang, Sylke Winkler, Frédéric Legendre, Dino P. McMahon, Mark C. Harrison

## Abstract

Subsociality and wood-eating or xylophagy are understood as key drivers in the evolution of eusociality in Blattodea (cockroaches and termites), two features observed in the cockroach genus *Cryptocercus*, the sister group of all termites. We present and analyse two new high-quality genomes from this genus, *C. punctulatus* from North America and *C. meridianus* from Southeast Asia, to explore the evolutionary transitions to xylophagy and subsociality within Blattodea. Our analyses reveal evidence of relaxed selection in both *Cryptocercus* and termites, indicating that a reduction in effective population size may have occurred in their subsocial ancestors. These findings challenge the expected positive correlation between dN/dS ratios and social complexity, as *Cryptocercus* exhibits elevated dN/dS values that may exceed those of eusocial termites. Additionally, we identify positive selection on mitochondrial ribosomal proteins and components of the NADH dehydrogenase complex, suggesting significant evolutionary changes in energy production. Future studies incorporating additional genomic data from diverse blattodean species are essential to elucidate the molecular mechanisms driving transitions to xylophagy and eusociality.

## Introduction

Subsociality, in which one or both parents care for the brood (Wilson, 1971; Tallamy & Wood, 1986), and xylophagy, the ability to feed on wood, have evolved multiple times within cockroaches (Blattodea) (Legendre & Grandcolas, 2018). For example, subsociality has emerged several times within the Blaberidae family, while xylophagy is found in the Nyctiboridae, Blaberidae, and Tryonicidae (Nalepa & Bell, 1997; Grandcolas, 1993). These two traits sometimes co-occur, at least within the genus *Salganea* (Blaberidae) and in *Cryptocercus* (Cryptocercidae) (Nalepa & Bell, 1997). The transition to eusociality, on the other hand, which involves alloparental care by non-reproducing relatives, has only occurred once in Blattodea: the termites (Termitoidae). Since *Cryptocercus* is the sister group to all termites (Lo et al., 2000; Inward et al., 2007; Legendre et al., 2008), it has been suggested that wood-feeding cockroaches that provide biparental care to their offspring represents the ancestral state of termites (Nalepa, 2010). Obtaining genomic data for *Cryptocercus* is therefore crucial for studying the molecular evolution of subsociality and xylophagy in cockroaches, as well as for understanding the circumstances under which the transition from subsociality to eusociality occurred within Blattodea (Legendre & Grandcolas, 2018).

Transitions to xylophagy involve complex adaptations necessary for the effective digestion of wood (Dillard & Benbow, 2020). This dietary shift frequently requires specific microbial communities and a supportive host gut environment for newly acquired (typically obligate) beneficial symbionts (Engel & Moran, 2013). In contrast to other cockroaches, both lower termites and *Cryptocercus* have obligate protist endosymbionts that are transferred to offspring via anal trophallaxis (Klass et al., 2008; Brune & Dietrich, 2015). Investigating how this dependence on gut symbionts has shaped the genomes of *Cryptocercus* species would provide valuable insight into the evolution of termite-microbial coexistence.

Furthermore, comparing the genomes of *Cryptocercus* with those of eusocial termites and other cockroaches is necessary to investigate the molecular signatures related to the emergence of subsociality and how they differ from mechanisms related to more complex social phenotypes. Previous studies have found sophisticaed chemical communication to be important for the evolution of eusociality in Hymenoptera, evidenced by high genomic copy numbers of odorant receptors (Kapheim et al., 2015; Shell et al., 2021; Simola et al., 2013), although this relationship between sociality and odorant receptor repertoires has been disputed (Gautam et al., 2024)). In termites, on the other hand, ionotropic receptors were predicted to be important for colony communication due to their caste-specific expression profiles, signals of positive selection, and high copy number, albeit reduced compared to the German cockroach, *Blattella germanica* (Harrison et al., 2018). Furthermore, in contrast to eusocial evolution in Hymenoptera (Shell et al., 2021), termite proteomes reveal significant gene family contractions compared with *B. germanica* (Harrison et al., 2018) and reductions in regulatory complexity (Jones et al., 2024). This is also accompanied by an overall reduction in genome size (Harrison et al., 2018; Terrapon et al., 2014). Across many origins of eusociality, major genomic changes in transcriptional regulation have been detected compared to non-eusocial relatives (Simola et al., 2013; Shell et al., 2021; Kapheim et al., 2015; Harrison et al., 2018). However, shifts in expression patterns are also likely to facilitate the evolution of parental care in subsocial species (Rehan & Toth, 2015; Johnson & Linksvayer, 2010; Thorne, 1997), suggesting that the aforementioned regulatory changes may have undergone evolutionary changes prior to the evolution of eusociality.

Eusociality has been linked to increased dN/dS due to relaxed selection possibly caused by a reduction in the effective population size (Chak et al., 2021; Weyna & Romiguier, 2021; Roux et al., 2024). Relaxed selection has been associated with eusociality in Hymenoptera (Weyna & Romiguier, 2021; Hunt et al., 2011) and termites (Ewart et al., 2024; Roux et al., 2024), indicating a possible universal link between eusociality and relaxed selection. Subsocial sister groups were, however, mostly lacking in these studies, so that it is currently unknown whether selection was already relaxed in subsocial ancestors. However, for a single species, *C. wrighti*, dN/dS values appeared intermediate between solitary cockroaches and eusocial termites (Roux et al., 2024).

The genomic signatures of sociality within the order Blattodea have so far been studied by comparing solitary or gregarious cockroaches with the social termites (Terrapon et al., 2014; Harrison et al., 2018; Mikhailova et al., 2023). The termite genomes revealed changes in transcriptional regulation, chemical communication and patterns of selection as well as a reductions in genome and proteome sizes (Mikhailova et al., 2023). Not including *Cryptocercus* means that these studies cannot determine whether genomic changes occurred at or before the emergence of termites, limiting the ability to investigate the precise relationship between genomic traits and the transition to advanced sociality in termites. To understand the genomic changes that predated the emergence of termites, we present high-quality genomes of two subsocial *Cryptocercus* cockroaches, *C. punctulatus* and *C. meridianus*, which provide a valuable new resource for studying the evolution of sociality and xylophagy in Blattodea.

## Results

### ***Cryptocercus*** genomes and proteomes more similar to termites than cockroaches

We *de novo* assembled a high-quality genome of the wood-roach *Cryptocercus punctulatus* using a combination of long ONT and short TELL-seq reads together with Hi-C data. Here we analyse this genome along with a second new *Cryptocercus* genome, *C. meridianus*, presented in a parallel study (Liu et al., 2025), which was sequenced and assembled using similar methods. High contiguity (N50 = 46 and 70 Mb, L50 = 10 and 6), as well as high completeness (Complete BUSCO = 99.5% and 99.7%), were achieved for both genomes, respectively (Table 1). These genome assemblies are more contiguous compared to other available blattodean assemblies: the highest N50 is 16 Mb for the termite *Macrotermes belicosus* (Qiu et al., 2023).

**Table 1:** Genome assembly information and statistics.

| Species | Coverage | Length | # contigs | N50 | L50 | BUSCO |
| --- | --- | --- | --- | --- | --- | --- |
| *C. meridianus* | 56x | 820 Mb | 1288 | 70.22 Mb | 6 | C:99.5%[S:98.1%,D:1.4%],F:0.3%,M:0.2%,n:1367 |
| *C. punctulatus* | 24x | 1189 Mb | 1490 | 45.68 Mb | 10 | C:99.7%[S:97.8%,D:1.9%],F:0.1%,M:0.2%,n:1367 |

At 1.2 Gb and 820 Mb, these *Cryptocercus* genome assemblies are more similar in size to those of six published termite genomes (median 0.9 Gb) than to those of other published cockroach genomes (median 3 Gb) (Fig. 1). Similarly, the proteomes of *Cryptocercus* (18,688 protein-coding genes in *C. punctulatus* and 15,909 in *C. meridianus*) and termites (from 12,984 in *Coptotermes formosanus* to 18,160 in *Cryptotermes secundus*) are substantially smaller than those of the other three analysed published cockroach genomes (*Blattella germanica* : 25 393, *Diploptera punctata* : 27 939, *Periplaneta americana* : 27 047; Fig. 1). Taken together, these findings indicate that a reduction in genome and proteome size occurred in a common ancestor of termites and *Cryptocercus*, along with or predating the evolution of subsociality and xylophagy.

**Figure 1:**
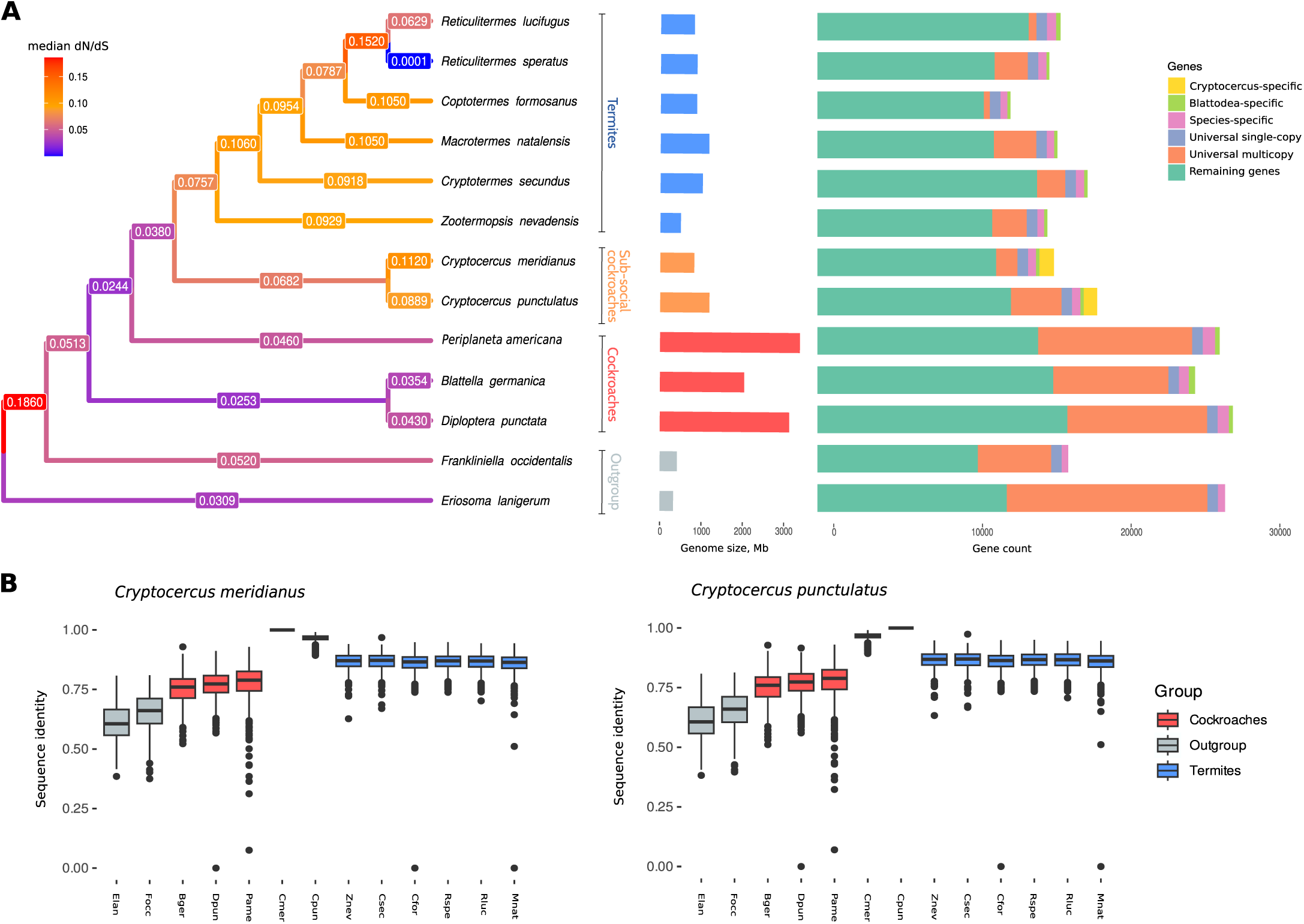
A: Phylogenetic tree of species used in this study with the colour showing median dN/dS on each branch; the asterisks mark *Cryptocercus* branches that were used as foreground in the selection analyses; the barplots represent genome size and gene counts for each species. B: Boxplots of sequence identity between *Cryptocercus* and other species; the sequence identity was calculated as a frequency of identical sites in multiple sequence alignments for each single-copy orthogroup.

In our species set of six termites, five cockroaches and two outgroups, we identified a total of 19,999 gene ortholog groups, including 712 single-copy orthologs (SCO) and 1067 *Cryptocercus*-specific ortholog groups. A further 1109 and 1610 genes were species-specific, singletons in *C. meridianus* and *C. punctulatus*, respectively. Interestingly, within the 712 SCOs, we found higher sequence similarity between *Cryptocercus* and termites (0.862 to 0.872 median sequence identity to *C. punctulatus* or *C. meridianus*) than with other cockroach species (0.760 to 0.789; Fig. 1B). As expected, sequence identity was high between *C. meridianus* and *C. punctulatus* (0.967). The loss in sequence identity to orthologs in *Cryptocercus* differs between termites and cockroaches, even when considering estimated divergence times. While orthologs in termites diverged from *Cryptocercus* species at a median of 0.070-0.076 percentage points per million years (pp/MY), they diverged at a median of 0.103 to 0.112 pp/MY in cockroaches.

### Relaxed selection with the evolution of subsociality

Previous studies have found a relaxation in selection strength in eusocial species due to an expected decrease in effective population size (N*_e_*) compared to non-eusocial species (Ewart et al., 2024; Roux et al., 2024). To test whether such a relaxation in selection could be detected as early as the onset of subsociality in *Cryptocercus*, we analysed dN/dS among all SCOs. We found generally higher values in *Cryptocercus* (mean: 0.140 & 0.164; median: 0.089 & 0.112) and termites (mean: 0.113-0.230; median: 0.063-0.105, except on the very short *R. speratus* branch: 0.0001) than in other non-subsocial, non-wood-feeding cockroaches (mean: 0.0481-0.0638; median: 0.035-0.046), indicating that relaxation in selection may have already occurred with the onset of subsociality (Fig. 1A). If increasing social complexity does indeed cause a relaxation of selection via a decrease in N*_e_*, then we would expect a stepwise increase in dN/dS in the three groups: 1) non-subsocial cockroaches; 2) subsocial *Cryptocercus*; 3) eusocial termites. To test this, we estimated dN/dS for different branch models with CodeML (Yang, 2007), in which we allowed a) a single dN/dS for all Blattodea (H0), b) two dN/dS values, one each in all cockroaches and in termites (H2a), c) or one each in non-social cockroaches and in social Blattodea (*Cryptocercus* plus termites; H2b), as well as d) three values, testing for differences in dN/dS among non-subsocial cockroaches, *Cryptocercus* and termites (H3). Model H2b was favoured over H0 most often (618 out of 712 SCOs at FDR *<* 0.05), followed by H3 (614) and H2a (581), indicating an increase in dN/dS occurred throughout *Cryptocercus* and termite evolution. Interestingly, when using the H3 model, dN/dS was significantly higher on *Cryptocercus* branches than on termite branches (W = 206793, p = 8.9x10*^−^*^10^), however, this model was only preferred over the H2b model for 144 out of the 712 orthologs. These findings support a potential relaxation of selection throughout termites and *Cryptocercus* but the data do not, with the current data set, support a link between relaxed selection and increasing social complexity.

In further support of increased dN/dS levels in *Cryptocercus* using RELAX from the HyPhy suite, we identified 18 of the 712 SCOs to have undergone a significant relaxation of selection on *Cryptocercus* branches (adjusted p-value *<* 0.05). These genes with significant signatures of relaxed selection contained 5 genes with mitochondrial functions, including two mitochondrial ribosomal proteins and an NADH dehydrogenase subunit of the electron transport chain (Table S1). Accordingly, these genes were enriched for the GO-term “mitochondrial translation”, as well as phosphatidyl phosphate biosynthesis, tRNA processing, primary metabolic process, and organic substance metabolic process (Fig. 2). We found only 5 genes undergoing significant intensified selection on *Cryptocercus* branches (adjusted p-value *<* 0.05) that interestingly also included two genes with mitochondrial functions, as well as two genes involved in RNA regulation and a component of the proteasome (Table S1).

**Figure 2:**
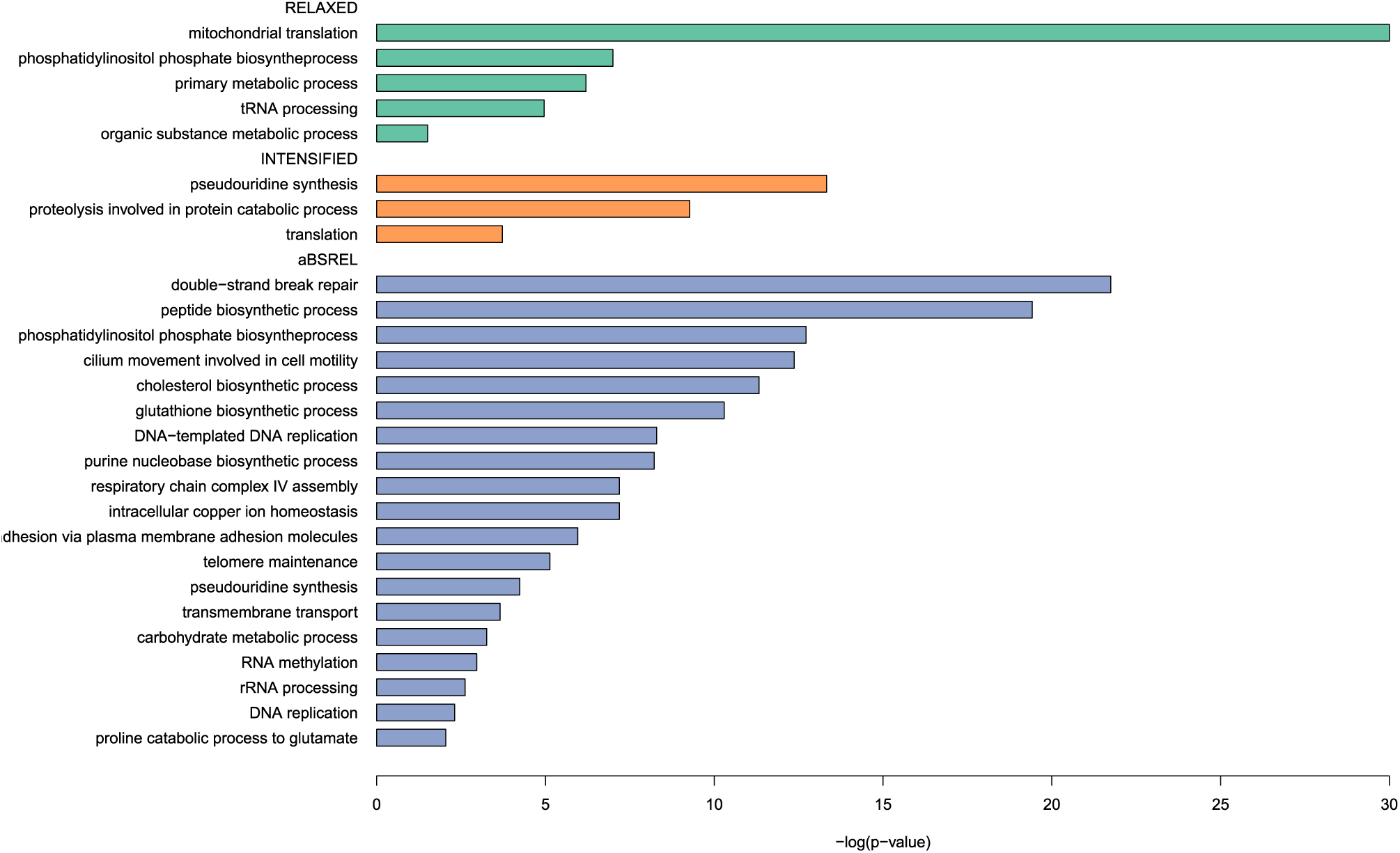
GO-terms significantly enriched in orthologs under relaxed, intensified or positive selection on *Cryptocercus* branches.

### Adaptive evolution in *Cryptocercus*

Besides signals of relaxed selection, we also found evidence for positive selection in *Cryptocercus*. With aBSREL from the HyPhy suite, we identified 63 SCOs that underwent positive selection in at least one of the three tested *Cryptocercus* branches (marked with ’*’ in Fig. 1A). We found four of these 63 genes to be under positive selection on two out of the three branches and a single gene on all three *Cryptocerus* branches. The gene under positive selection on all three *Cryptocercus* branches (OG0005583, Cmer00006637, Cpun00005036) is predicted to be the mitochondrial ribosomal protein, L47. Interestingly, another ortholog under positive selection on both terminal branches (OG0004724, Cmer00001705, Cpun00017128) is also predicted to be a component of a mitochondrial ribosomal protein, S26. A further mitochondrial gene, NADH dehydrogenase PDSW subunit (ND-PDSW; OG0004337, Cmer00013000, Cpun00010840) was under positive selection on two *Cryptocercus* branches (the *C. punctulatus* branch and the ancestral branch). The other genes under positive selection on two *Cryptocercus* branches were orthologous to Lrch (Leucine-rich-repeats and calponin homology domain protein; OG0005620 both terminal branches) and CG1090 (involved in calcium homeostasis; OG0004846 on the *C. meridianus* branch and the ancestral branch). A further 7 components of the mitochondrial ribosome (mRpL15, mRpL17, mRpL22, mRpL28, mRpL9, mRpS27, mRpS30), as well as the 39 kDa subunit of NADH dehydrogenase were under significant positive selection on one of the three *Cryptocercus* branches. The 63 significant genes also include 15 additional genes with functions related to translational or transcriptional regulation. Overall, these 63 genes under significant positive selection were enriched for GO terms such as respiratory chain complex IV assembly, translation release factor activity, phosphatidylinositol phosphate biosynthesis, DNA repair, and RNA methylation (Fig. 2).

### Gene family evolution

We investigated the evolution of gene family size with CAFE5 (Mendes et al., 2020). In line with the observed reduction in proteome size, we inferred a substantially higher number of total gene family contractions than expansions at the ancestral node of *Cryptocercus* and termites (316 and 75, respectively), including 54 significant contractions and 33 significant expansions (Fig. 3). At the origin of *Cryptocercus*, we also inferred a higher number of contractions than expansions (546 & 230). However, a larger number of gene families were significantly expanded (63) rather than contracted (40) (Fig. 3). On each *Cryptocercus* terminal branch we found similar numbers of expansions and contractions, although expansions were slightly higher in *C. punctulatus* (expansions: 417, 199 significant; contractions: 308, 133 significant), while contractions were higher in *C. meridianus* (expansions: 272, 112 significant; contractions: 399, 175 significant; Fig. 3).

**Figure 3:**
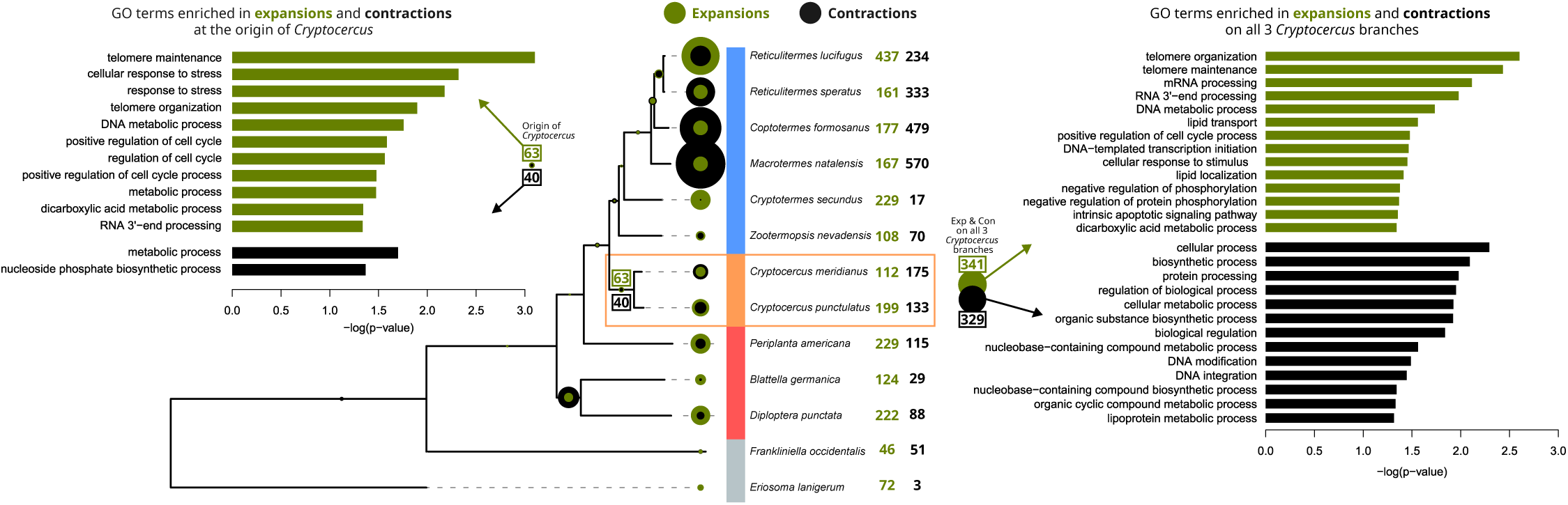
Gene family evolution. Shown are the numbers of significant gene family expansions (green) and contractions (black) throughout the analysed phylogeny. The bar plots show the GO-terms that are significantly enriched among expansions and contractions at the root of *Cryptocercus* (left) and on all 3 *Cryptocercus* branches (right).

Among those genes significantly expanded at the origin of *Cryptocercus*, we found a significant enrichment of 11 ’BP’ GO-terms, including functions related to telomere maintenance, DNA-repair and cell-cycle, as well as tricarboxylic acid cycle, autophagy of mitochondrion and 2-oxoglutarate metabolic process (Fig. 3). When considering all gene families expanded on any of the three *Cryptocercus* branches, we also found significantly enriched GO-terms related to mRNA processing, lipid transport, negative regulation of phosphorylation and intrinsic apoptotic signaling (Fig. 3).

The genes significantly contracted at the origin of *Cryptocercus* were enriched for metabolic process and nucleoside phosphate biosynthetic process (Fig. 3). Among gene families significantly contracted on any of the *Cryptocercus* branches, we also found enrichments of GO-terms related to biological regulation, protein processing and lipoprotein metabolism (Fig. 3).

### Ionotropic Receptors are reduced in *Cryptocercus* and termites

Previous studies suggest an expansion of Ionotropic Receptors (IRs) within Blattodea, with hundreds of copies found in the genomes of the German cockroach (Harrison et al., 2018), the American cockroach (S. Li et al., 2018) and the Pacific beetle cockroach (Fouks et al., 2023). However, these chemoreceptors were shown to be reduced to around 100 in three termite species (Harrison et al., 2018). Similar to previous findings, we predicted over 400 IR copies in each of the three analysed cockroach genomes (*D. punctata* : 434; *B. germanica* : 978; *P. americana*: 442) and fewer than half as many in five of the analysed termite species (106 to 213) (*R. lucifugus* was not analysed due to missing genomic information). In the *Cryptocercus* genomes we predicted 199 in *C. meridianus* and 201 gene copies in *C. punctulatus*, suggesting that the reduction in the IR repertoire predated the evolution of *Cryptocercus* and termites. In each of the two outgroups (two Acercaria quite distantly related to Blattodea), we predicted only 28 copies.

### Predicted methylation of genes

We predicted methylation levels of each gene for all of the analysed genomes by comparing CpG counts to expectations based on proportions of cytosines and guanines in coding regions. A depletion of CpGs is an indication of increased DNA methylation (Bewick et al., 2017). We found a unimodal distribution of CpG*_o/e_* values in the genes of the three non-social cockroaches but a bimodal distribution in *Cryptocercus spp.* and in all analysed termites, except *C. meridianus*, which may have 3 modes (Fig. 4). These results indicate that a subsection of genes are more strongly methylated than other genes in *Cryptocercus* and termites.

**Figure 4:**
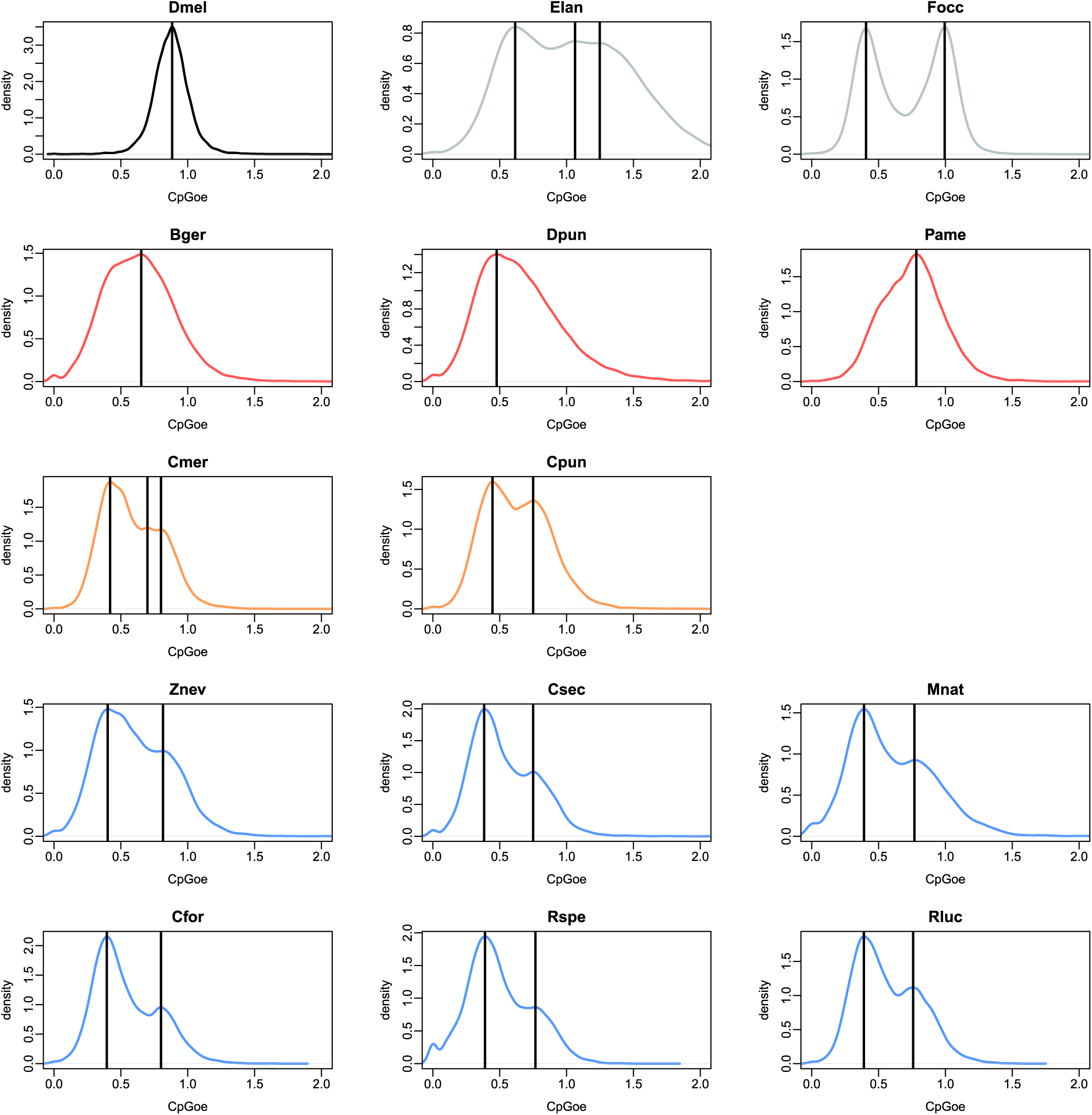
Distributions of gene CpG*_o/e_* values for the analysed species. Modes are denoted as vertical lines.

## Discussion

To investigate genomic changes during a social transition in Blattodea, we analysed two high-quality genomes of subsocial wood-feeding *Cryptocercus* cockroaches which represent the nearest extant sister lineage of termites. With these novel resources, we have been able to test whether previously reported genomic differences between termites and cockroaches are uniquely associated with the evolution of advanced sociality in termites, or whether they coincide with the emergence of wood-feeding and subsociality in the common ancestor of termites and *Crytocercus*. Confirming previous C-value estimations of genome size (Koshikawa et al., 2008), we found that a reduction of genome size likely occurred in the common ancestor of *Cryptocercus* and termites, either coinciding with or predating transitions to subsociality and wood-feeding. We also found smaller proteomes both within *Cryptocercus* and termites compared to non-subsocial cockroaches. This was exemplified by the number of ionotropic receptors (IRs) we detected in the analysed Blattodean genomes. IRs had previously been inferred to be massively expanded in cockroaches and then secondarily reduced in termites (Harrison et al., 2018; S. Li et al., 2018; Fouks et al., 2023). Our analyses support a reduction in IRs in both extant *Cryptocercus* and termite species compared to the three rather generalist non-subsocial cockroaches included in this study, indicating either that a reduced number of chemoreceptors is linked with more integrated forms of sociality and/or wood-feeding. Genome assemblies of other wood-feeding and omnivorous cockroaches and of other more distantly related subsocial cockroaches are necessary to investigate this question further.

We show that the previously predicted change in methylation patterns in termites compared to cockroaches (Harrison et al., 2018) may have evolved much earlier than previously thought: in the common ancestor of *Cryptocercus* and termites, and therefore coinciding with the presumed transition to subsociality. These results support the hypothesis that fundamental gene regulatory changes were necessary for the initial evolutionary transition to subsociality in *Cryptocercus* (Mikhailova et al., 2023), implying that this was a necessary genomic prerequisite for the evolution of advanced sociality in termites, given that such methylation patterns are conserved across termites. In further support, we present evidence for the adaptive evolution of several genes involved in translation and transcriptional regulation within the *Cryptocercus* lineage. Whether our proposed methylation patterns or indeed other mechanisms of transcriptional regulation changed to such a dramatic degree in the common ancestor of termites and *Cryptocercus* needs to be further corroborated with more direct measurements, such as bisulfite sequencing and transcriptomic sequencing across a broader range of blattodean species. Such data will help to illuminate the role of transcriptional regulation in the evolution of subsociality and its potential importance for the emergence of phenotypic plasticity in termites.

Additionally, we present evidence for a relaxation of selection in both *Cryptocercus* and termite genomes, indicating that a reduction in effective population size may already have occurred in the presumed subsocial, wood-feeding ancestor of this clade. The reasons for this apparent relaxation of selection in *Cryptocercus* and in termites require further investigation. Our results confirm previous findings of inflated dN/dS values in termites compared to cockroaches (Ewart et al., 2024; Roux et al., 2024) but importantly cast doubt on the expectation of a positive correlation between dN/dS and social complexity due to a hypothesised reduction in N*_e_* in more advanced social groups (Roux et al., 2024; Ewart et al., 2024; Mikhailova et al., 2023; Weyna & Romiguier, 2021). In contrast to a previous study based on transcriptomes, in which dN/dS values for *C. wrighti* were intermediate between termites and non-social cockroaches (Roux et al., 2024), in the current analysis using whole genomes, we find at least equally high and possibly higher dN/dS values in subsocial *Cryptocercus* compared to eusocial termites. Our findings suggest that relaxed selection was linked to the transition to wood-feeding and parental care in the common ancestor of termites and *Cryptocercus* rather than to the emergence of advanced sociality in termites. However, using current data, we are unable to separate the effects of phylogeny, wood-feeding and sociality on dN/dS levels. The integration of further higher quality genomes from species with varying levels of social complexity throughout Blattodea will allow this question to be more rigourously interrogated.

We found strong signatures of selection on mitochondrial functions, indicating a potential association between adaptive changes in ATP production and wood-feeding. Nine mitochondrial ribosomal proteins and two components of NADH dehydrogenase of the mitochondrial electron transport chain were under positive selection, while a further component each of NADH dehydrogenase and the mitochondrial ribosome experienced intensified selection on *Cryptocercus* branches. Interestingly, five genes with mitochondrial functions were also under strongly relaxed selection, including two mitochondrial ribosomal proteins and the SGDH component of NADH dehydrogenase (Table S1).

These two *Cryptocercus* genomes provide important genomic resources both for understanding social evolution and the transition to wood-feeding (Legendre & Grandcolas, 2018). We present evidence for genomic adaptations to feeding on wood; a recalcitrant and energy-poor resource. Here, we show that some genomic traits previously associated with termites, such as elevated dN/dS (Ewart et al., 2024) and global patterns of methylation (Harrison et al., 2018) likely evolved much earlier, at the earliest in the most recent common ancestor of *Cryptocercus* and termites. This suggests that the selective pressures linked to the emergence of subsociality, xylophagy as well as the acquisition of novel obligate gut symbionts (Biedermann & Rohlfs, 2017), and that understanding the evolutionary origins of termite sociality require a more detailed and focused analysis of gene family as well as gene regulatory evolution. Equally, we encourage increased focus on diversity across Blattodea. The generation of further high-quality genomes from species varying in degrees of (sub)sociality and diet, from species representing independent origins of xylophagy and subsociality, is essential for investigating the genomic mechanisms involved in the evolution of two major evolutionary novelties in this clade: xylophagy and eusociality.

## Methods

### Genome sequencing

#### Extraction of genomic DNA

*Cryptocercus punctulatus* individuals were collected in Virginia (USA) and reared in the ISYEB laboratory. High molecular weight genomic DNA (HMW gDNA) was extracted from snap-frozen single legs of *C. punctulatus* with the Circulomics Insect Big DNA kit (protocol version v0.20a). In brief, one leg was homogenised on ice for 10s with the TissueRuptor (Qiagen), and the homogenised tissue was lysed in buffer CT (Circulomics) with Proteinase K. RNA was removed by RNAse A treatment. HMW gDNA was captured with the Circulomics Nanodisks and eluted in elution buffer. This protocol resulted in high-quality HMW gDNA of about 170 Kb in size when analysed with the Agilent Femtopulse capillary electrophoresis system.

#### Oxford Nanopore (ONT) sequencing of HMW gDNA

One ONT library of HMW gDNA of *C. punctulatus* was prepared as recommended by Oxford Nanopore according to the Nanopore protocol ’Genomic DNA by ligation’ (SQK-LSK110, Version: GDE_9108_v110_revH_10Nov 2020). In brief, approximately 4 *µ*g of HMW gDNA was used for the ONT library preparation, after applying a DNA repair and end repair step, ONT sequencing adapters were ligated to the HMW gDNA. The final ONT library was loaded twice on a PromethION FLO-PRO002 flow cell with the reload after 24 hours and ran in total for 3 days on the PromethION instrument of the Vienna Biocenter Core Facilities. Sequencing yield was 33 Gbp, average length of the library about 7.5 kb.

#### TELL-Seq whole genome sequencing

One indexed paired-end Transposase Enzyme Linked Long-read Sequencing (TELL-Seq) whole genome library was prepared from 10 ng of HMW gDNA extracted from one snap-frozen single leg of *C. punctulatus* following the TELL-Seq WGS Library Prep User Guide (Document # 100005 v6). In brief, fragments of HMW gDNA were barcoded with unique molecular identifiers (= barcodes) making use of barcoded microbeads by a transpose. This barcoding step is followed by a second transposase step to fragment the barcoded gDNA and to add an indexing barcode. After amplification and further clean-up, the resulting barcoded linked-read TELL-Seq library ran on the NovaSeq 6000 of the DRESDEN concept genome center on an S4 flow cell with 2x 150 cycles.

#### Hi-C confirmation capture

Chromatin confirmation capturing was done making use of the ARIMA-HiC 2.0 kit and followed the user guide for animal tissues (ARIMA Document, Part Number: A160162 v00). In brief, 1 flash-frozen powdered leg of *C. punctulatus* was cross-linked chemically. 760 ng cross-linked genomic DNA was digested with the ARIMA-HiC 2.0 restriction enzyme cocktail consisting of four restriction enzymes. The 5â-overhangs were filled in and labelled with biotin. Spatially proximal digested DNA ends were ligated and finally the ligated biotin containing fragments were enriched and went for Illumina library preparation, which followed the ARIMA user guide for Library preparation using the KapaR Hyper Prep kit (ARIMA Document Part Number A160139 v00). The barcoded Hi-C library ran on an S4 flow cell of a NovaSeq6000 with 2x 150 cycles.

### Assembly

The assembly of *C. punctulatus* genome was performed as described in a previous study (Liu et al., 2025). In short, the long ONT reads were assembled using Flye version 2.9 with default settings (Kolmogorov et al., 2019). The quality of the assembly was evaluated using QUAST version 5.0.2 (Mikheenko et al., 2018) and BUSCO version 5.1.2 (Manni et al., 2021) after each assembly step. The assembly was polished with short TELL-seq reads using HyPo version 1.0.3 (Kundu et al., 2019) with default settings. To resolve regions with high heterozygosity, we used purge haplotigs (Roach et al., 2018). Initially, a coverage histogram was created using the command “purge_haplotigs hist”. Subsequently, coverage was analyzed for each contig individually with the command “purge_haplotigs cov -l 5 -m 70 -h 250” with the argument values chosen based on the coverage histogram. The scaffolding was done using Hi-C data and the following programs: mapped Hi-C reads with BWA-MEM (H. Li, 2013), processed with pairtools (Open2C et al., 2024) with the command “pairtools sort –nproc 16”. PCR and optical duplicates were removed with the command “pairtools dedup –nproc-in 8 –nproc-out 8 –mark-dups”. The mapped reads were used to generate scaffolded genome assembly with YaHS version 1.2a.2 with default settings (Zhou et al., 2023).

### Annotation

The *C. punctulatus* genome was annotated using the same pipeline as for *C. meridianus* (Liu et al., 2025). First, repeat elements were detected by RepeatModeler v.2.0.3 (Smit et al., 2015b) and masked by RepeatMasker v.4.1.2 (Smit et al., 2015a). The masked genome was used for identifying protein-coding genes, which integrated protein-genome alignments, transcripts-genome alignments and *ab initio* gene predictions. For protein-genome alignments, we collected protein sequences of (1) 45 termite species, *C. meridianus* and *Blatta orientalis* Liu et al. (2025), (2) 10 model insects (*Aedes aegypti*, *Anopheles gambiae*, *Drosophila melanogaster*, *Acyrthosiphon pisum*, *Apis mellifera*, *Nasonia vitripennis*, *Solenopsis invicta*, *Bombyx mori*, *Manduca sexta*, *Tribolium castaneum*) from NCBI, and (3) four Blattodeans (*Blattella germanica*, *Cryptotermes secundus*, *Zootermopsis nevadensis*, *Coptotermes formosanus*) from InsectBase (Mei et al., 2022), and mapped them to the genome using miniprot v.0.7 (H. Li, 2023). For transcript-genome alignments, RNA reads were mapped to the masked genome using Hisat2 v.2.2.1 (Kim et al., 2019) and assembled using StringTie v.2.1.4 (M. Pertea et al., 2015). As for *ab initio* predictions, we incorporated (1) AUGUSTUS v.3.4.0 (Stanke & Waack, 2003) trained with gene structures identified from transcript-genome alignments by TransDecoder v.5.6.0 (B. Haas et al., 2016), (2) BRAKER v.2.1.6 (Hoff et al., 2019) trained with RNA read-genome mapping from Hisat2, (3) GALBA v.1.0.1 (Bruna et al., 2023) trained with protein sequences of the four Blattodeans from InsectBase. Then protein-genome alignments, transcripts-genome alignments, gene structures from TransDecoder and *ab initio* gene predictions were integrated into consensus gene structures by EvidenceModeler v.1.1.1 (B. J. Haas et al., 2008). Genes structures were removed if they were inferred by only one *ab initio* predictor and lack additional evidence from other sources. These gene structures and transcripts assembled by StringTie were processed with two iterations of PASA v.2.5.2 (B. J. Haas et al., 2008). Genes were excluded if they have incomplete reading frame or in-frame stop codon. Finally, we extracted the longest protein isoform of each gene, searched them against the non-redundant (nr) database using DIAMOND v2.1.7.161 (Buchfink et al., 2015), and processed the homology hits using MEGAN6 (Huson et al., 2007), which assigned peptides to taxa. Scaffolds were excluded as contaminations if above 90% of genes on them were assigned to Bacteria, Archaea or viruses.

### Preparation of proteomes for analyses

For further analyses, we added the existing genomic data of 3 cockroaches and 6 termites as well as two outgroup species (*Eriosoma lanigerum* and *Frankliniella occidentalis*) (Harrison et al., 2018; S. Li et al., 2018; Fouks et al., 2023; Terrapon et al., 2014; Shigenobu et al., 2022; Itakura et al., 2020; Poulsen et al., 2014; Biello et al., 2021; Rotenberg et al., 2020; Martelossi et al., 2023). We extracted the proteomes with gffread (G. Pertea & Pertea, 2020), then kept only the longest isoforms and filtered out pseudogenes using DW-Helper scripts (https://domain-world.zivgitlabpages.uni-muenster.de/dw-helper/).

### Gene dynamics and selection analysis

To investigate selection and gene family dynamics we identified orthologs using orthofinder2 (Emms & Kelly, 2018). Gene family dynamics were investigated using CAFE5 with a single K and lambda value having the highest maximum likelihood (Mendes et al., 2020). Protein-guided multiple sequence alignments were run using PRANK (Löytynoja, 2014) and GUIDANCE (Penn et al., 2010) and trimmed using Gblocks (Talavera & Castresana, 2007). Overall dN/dS ratios were found using codeml (Yang, 2007) while branch-specific relaxed selection was identified using RELAX (Wertheim et al., 2015) and branch-specific selection using aBSREL (Smith et al., 2015). Significant results were determined with a p-value below 0.05 for aBsrel and at the same cut-off after FDR correction for RELAX. Protein domain scans were carried out using Pfamscan (Finn et al., 2014) and interproscan (Mitchell et al., 2015) with GO terms extracted using Pfam2GO. Individual GO universes were created for both selection and CAFE analyses, while enrichment was conducted with TopGO (Alexa & Rahnenführer, 2009).

### Gene divergence

Sequence identity of *Cryptocercus* genes compared to other analysed blattodean genomes was calculated based on the alignments of single-copy orthologs from the previous step. We used “seqidentity” function from the R package bio3d to calculate the percentage of identical sites within each orthologous group (Grant et al., 2006). To calculate relative differences in sequences, we divided loss of identity (1 - sequence identity) by divergence times between pairs of species. Divergence times were taken from timetree.org (accessed September 2024) based on (Bourguignon et al., 2018) for most species and (Forni et al., 2021) for dating the Blaberoidea species *B. germanica* and *D. punctata*.

### Predicted methylation profiles

Relative levels of CpG depletion were calculated per gene using a custom Python script. In brief, for each gene, the number of observed CGs in the coding region was divided by the expected number of CGs (CpG*_e_*). CpG*_e_* is calculated from the the product of the individual proportions of Cs and Gs in the sequence.

### Annotation of ionotropic receptors

Ionotropic receptors were annotated in the analysed genomes with Bitacora, version 1.4 (Vizueta et al., 2020). Protein sequences of known IRs were taken from (Fouks et al., 2023) and aligned with mafft, v. 7.505 (Katoh et al., 2002), with the settings: –genafpair –maxiterate 16. A hidden markov model was created from the alignment with hmmbuild from the hmmer suite, v. 3.3 (Mistry et al., 2013). Bitacora was run on this hmm and all annotated protein sequences for each species, using an e-value threshold of 1e-05, including GeMoMa, v. 1.7.1 (Keilwagen et al., 2019), a maximum intron length of 15 000, and a filter length of 30 amino acids.

## Acknowledgments

ARCJ acknowledges support from the DFG through grant BO 2544 / 15-1 to E.B.-B. This work was supported by French National Research Agency grant no ANR-19-CE02-0023 (project Sociogenomics) to FL. AM was supported by the DFG grant HA 8997/1-1 to MCH. CA was supported by the DFG grant MC 436/5-1 to DPM. We are grateful to the following members of the DcGC Dresden-concept Genome Center for their contributions to genome sequencing: Nicola Gscheidel (Hi-C and TELL-Seq experiments), Wenhua Tan (ONT library preparation and sequencing), and Montserrat Palau de Miguel (optimisation of gDNA extraction).

## SUPPLEMENTARY MATERIAL

### Supplementary figures

**Figure S1:**
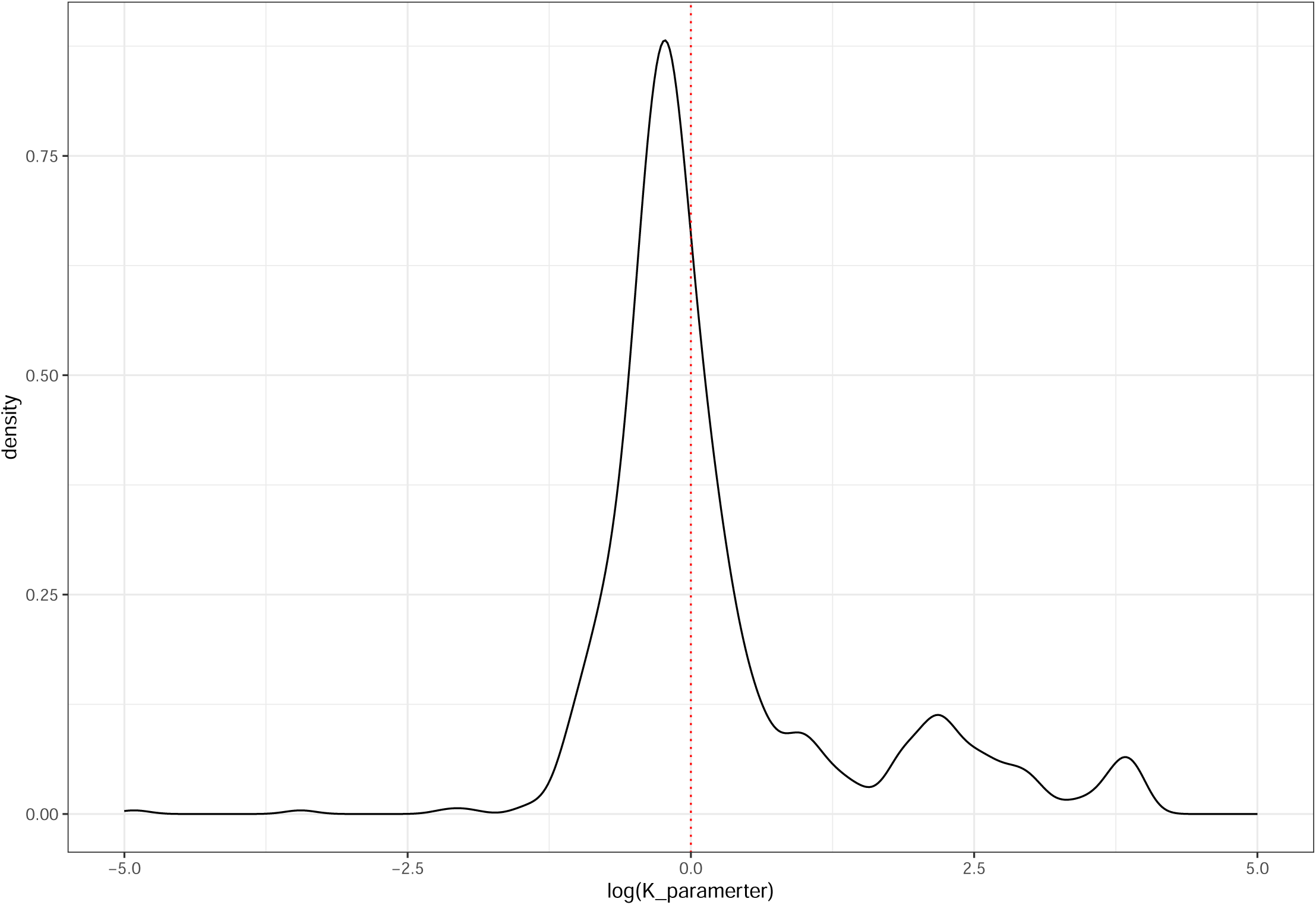
Density plot of RELAX K parameter estimates

### Supplementary figures

**Table S1:** Single-copy orthologues with significant signals of relaxed and intensified selection in *Cryptocercus* with the corresponding orthologue in *Drosophila melanogaster*.

| Orthogroups | K value | adjusted P | *C. punctulatus* | *C. meridianus* | *D. melanogaster* | Description |
| --- | --- | --- | --- | --- | --- | --- |
| RELAXED |  |  |  |  |  |  |
| 1. OG0005418 | 0.007 | $2.9 \times 10^{-4}$ | Cmer00010420-R0_1 | Cpun00007014-R0_1 | mRpL9 | Structural constituent of ribosome, involved in mitochondrial translation (mitochondrial ribosomal protein L9). |
| 2. OG0005006 | 0.139 | 0.005 | Cmer00014627-R0_1 | Cpun00014592-R0_1 | Notum | Cleaves glycoposphatidylinositol anchors |
| 3. OG0005210 | 0.234 | 0.011 | Cmer00007087-R0_1 | Cpun00000310-R1_1 | mRpS35 | Part of mitochondrial small ribosomal subunit, orthologous to human MRPS35 (mitochondrial ribosomal protein S35). |
| 4. OG0004879 | 0.308 | $7.7 \times 10^{-4}$ | Cmer00001706-R0_1 | Cpun00017126-R0_1 | CG4679 | Mitochondrial translation |
| 5. OG0005282 | 0.354 | $2.4 \times 10^{-4}$ | Cmer00002860-R1_1 | Cpun00010651-R2_1 | nonA | Required for normal vision and courtship behavior in *Drosophila*. |
| 6. OG0005473 | 0.360 | $3.4 \times 10^{-6}$ | Cmer00006416-R0_1 | Cpun00004667-R0_1 | Pi3K59F | Phosphatidylinositol 3 kinase 59F (Pi3K59F), required for formation of Phosphatidylinositol 3-phosphate. |
| 7. OG0005378 | 0.368 | $6.2 \times 10^{-4}$ | Cmer00011206-R1_1 | Cpun00007691-R0_1 | Dus2 | tRNA-dihydrouridine20 synthase activity. |
| 8. OG0004935 | 0.373 | 0.003 | Cmer00002694-R0_1 | Cpun00002791-R1_1 | CG2260 | Maturation of SSU-rRNA from tricistronic rRNA transcript |
| 9. OG0005690 | 0.381 | 0.004 | Cmer00007913-R0_1 | Cpun00015846-R0_1 | m | Belongs to the ZP-domain proteins family known as extracellular modular proteins. |
| 10. OG0005474 | 0.387 | 0.027 | Cmer00000965-R0_1 | Cpun00008630-R0_1 | Mccc2 | Carboxyltransferase subunit of the 3-methylcrotonyl-CoA carboxylase |
| 11. OG0005598 | 0.407 | 0.043 | Cmer00005847-R0_1 | Cpun00009826-R0_1 | Sur-8 | MAPK pathway activation |
| 12. OG0005642 | 0.458 | 0.043 | Cmer00001753-R0_1 | Cpun00014948-R1_1 | ND-SGDH | NADH dehydrogenase SGDH subunit; mitochondrial electron transport chain |
| 13. OG0004953 | 0.467 | 0.001 | Cmer00008734-R1_1 | Cpun00002549-R0_1 | ALIX | Adaptor protein implicated in multiple cellular processes |
| 14. OG0004518 | 0.472 | 0.001 | Cmer00000979-R0_1 | Cpun00008645-R0_1 | YME1L | AAA protease involved in mitochondrial quality control |
| 15. OG0004708 | 0.534 | 0.018 | Cmer00008444-R2_1 | Cpun00003086-R1_1 | kuz | ADAM metalloendopeptidase |
| 16. OG0005637 | 0.548 | 0.012 | Cmer00002733-R1_1 | Cpun00002759-R0_1 | pix | ATPase that functions as a translation recycling factor |
| 17. OG0005654 | 0.552 | 0.027 | Cmer00010593-R0_1 | Cpun00006597-R0_1 | Ddx1 | Member of the DEAD box family of RNA helicases |
| 18. OG0005051 | 0.585 | 0.012 | Cmer00004216-R0_1 | Cpun00009319-R0_1 | fw | Selectin that mediates the interaction of planar cell polarity |
| INTENSIFIED |  |  |  |  |  |  |
| 1. OG0004945 | 50 | 0.050 | Cmer00001139-R0_1 | Cpun00016824-R0_1 | Prosbeta6 | Non-catalytic component of the proteasome |
| 2. OG0005542 | 7.888 | $3.4 \times 10^{-6}$ | Cmer00010398-R0_1 | Cpun00006694-R0_1 | Pus1 | Pseudouridine synthase activity |
| 3. OG0004337 | 5.334 | 0.012 | Cmer00013000-R0_1 | Cpun00010840-R0_1 | ND-PDSW | NADH dehydrogenase PDSW subunit; mitochondrial electron transport chain |
| 4. OG0005583 | 2.469 | 0.036 | Cmer00006637-R0_1 | Cpun00005036-R0_1 | mRpL47 | Component of mitochondrial ribosome, orthologous to human MRPL47 |
| 5. OG0005325 | 2.369 | 0.038 | Cmer00000889-R0_1 | Cpun00008552-R0_1 | CG4806 | mRNA binding activity |

